# Coupled abiotic-biotic cycling of nitrous oxide in tropical peatlands

**DOI:** 10.1101/2022.01.14.475290

**Authors:** Steffen Buessecker, Analissa F. Sarno, Mark C. Reynolds, Ramani Chavan, Jin Park, Marc Fontánez Ortiz, Ana G. Pérez-Castillo, Grober Panduro Pisco, José David Urquiza-Muñoz, Leonardo P. Reis, Jefferson Ferreira-Ferreira, Jair M. Furtunato Maia, Keith E. Holbert, C. Ryan Penton, Sharon J. Hall, Hasand Gandhi, Iola G. Boëchat, Björn Gücker, Nathaniel E. Ostrom, Hinsby Cadillo-Quiroz

## Abstract

Atmospheric nitrous oxide (N_2_O) is a potent greenhouse gas thought to be mainly derived from microbial metabolism as part of the denitrification pathway. Here, we report that in unexplored peat soils of Central and South America, N_2_O production can be driven by abiotic reactions (≤ 98 %) highly competitive to their enzymatic counterparts. Extracted soil iron positively correlated with *in-situ* abiotic N_2_O production determined by isotopic tracers. Moreover, we found that microbial N_2_O reduction accompanied abiotic production, essentially closing a coupled abiotic-biotic N_2_O cycle. Anaerobic N_2_O consumption occurred ubiquitously (pH 6.4-3.7), with proportions of diverse clade II N_2_O-reducers increasing with consumption rates. Our findings show denitrification in tropical peat soils is not a purely biological process, but rather a “mosaic” of abiotic and biotic reduction reactions. We predict hydrological and temperature fluctuations differentially affect abiotic and biotic drivers and further contribute to the high N_2_O flux variation in the region.

## Introduction

The atmospheric accumulation of nitrous oxide (N_2_O), a potent greenhouse gas, has continued to increase^1,2^, calling for a better mechanistic understanding of its sources and sinks. Tropical soils are a major source of N_2_O. The largest contribution to global N_2_O flux, along with the highest uncertainties, have been observed over South America^3–5^ with large flux variations described in ground-based measurements from extensive peatlands of the Amazon basin^6,7^.

In waterlogged tropical peat soils, anoxic, reducing, humic acid-rich, and Fe-holding conditions are favorable for the abiotic formation of N_2_O^8^. Nitrous oxide can abiotically form from the reduction of nitrite (NO_2_^-^) via intermediary nitric oxide (NO), or hydroxylamine (NH_2_OH)^9^, both of which have typically low steady-state concentrations in soils. Hydroxylamine conversion into N_2_O relies on oxidants such as manganese (IV) minerals that are unlikely to persist in sufficient levels in the reducing milieu of peat. Thus, peatlands would generally favor the spontaneous chemical reduction of nitrogenous compounds – also called chemodenitrification. Some environments appear to sustain abiotic N_2_O production rates based on dissolved Fe and Fe mineral phases^10–12^, while others have shown an influence from organic matter (OM)^8^, presumably by providing complexed Fe^2+^ and/or humic electron shuttles^13^. Abiotic N_2_O formation has been recorded in polar^10,14^ and temperate^11^ environments, but the extent and distribution of this process in tropical peatlands have remained unexplored. With a recently estimated area of 1.7 million km^2^ (ref. ^15^), tropical peatlands under varying climatic regimes could play a major role in global N_2_O gas cycling.

Denitrification, generally occurring at oxygen concentrations below 6 μM^16^, is considered to be driven predominantly by microbial communities using Fe- and Cu-dependent reductase enzymes^17^ through a modular pathway structure with different populations mediating only one or two reduction steps^18,19^. Denitrifying microbes are well adapted to the conditions found in peat soils because they anaerobically respire organic substrates using nitrogen oxides as terminal electron acceptors^20,21^. Also, the extensive N_2_O sink potential previously observed in diverse soils^22,23^ can be better explained with the discovery of the abundant clade II N_2_O-reducing bacteria. While clade I N_2_O-reducers are affiliated to the *Proteobacteria,* clade II N_2_O reducers are more diverse and scattered across multiple phyla^24^. Interestingly, the clade II members tend to lack NO_2_^-^ reductases more so than clade I members^24^. From an ecological perspective, this trait might correspond with an intrinsic capability of the soil habitat to reduce NO_2_^-^ via chemodenitrification. Cellular resources can be saved and relocated to the expression of NO and N_2_O reductases^25^ to catalyze a thermodynamically more favorable redox reaction (*ΔG* of N_2_O reduction is ~100 kJ mol^−1^ higher than *ΔG* of NO_2_^-^ reduction).

While interactions between microbial guilds have been proposed as the basis for modularity^23^, the interplay of denitrifiers with abiotic reactions has received little attention, even though chemodenitrification can reduce or contribute to different inorganic nitrogen pools, including N_2_O and NO. The compatibility of abiotic N_2_O production and modular microbial denitrification led us to hypothesize that a coupled abiotic-biotic N_2_O cycle could operate in tropical peatlands. To test our hypothesis, we explored the dynamics and underlying factors of abiotic N_2_O formation and microbial N_2_O reduction in six peatlands located across Central and South America using isotopic tracers. Simultaneously, we quantified and sequenced the *nosZ* gene as a marker for the N_2_O-reducing microbial community. Our results provide evidence for concomitant abiotic N_2_O production and microbial consumption active under various peat soil conditions.

## Results

### Fe^2+^ drives abiotic formation of N_2_O in high-N_2_O soils

We assessed soil denitrification in six tropical peatlands, of which four were located within or near the Amazon basin (San Jorge, SJO; Melendez, MEL; Sítio do Cacau, SCB, Fazenda Córrego da Areia, FCA) and two in Central America (Medio Queso, MQE and Las Vueltas, VUL) (Fig. 1). Measured steady-state concentrations of NO_2_ in soil pore water were below detection (< 1 μM) in the majority of sites, indicating rapid cycling^26,27^. The iron content and redox balance were highly variable, with higher Fe^2+^ concentrations in mountainous peat (~5 mM) and lower Fe^2+^ in oligotrophic peat (0.01-0.04 mM). To determine abiotic N_2_O production rates under near-natural conditions, we induced a ten-fold spike with ^15^NO_2_^-^ *in-situ* and measured ^14^N^15^NO + ^15^N^14^NO + ^15^N^15^NO evolution. Biotic activity was arrested by amending the soil with 87.5 mM ZnCl_2_. Because the addition of Zn can liberate Fe^2+^ ions inevitably stimulating N_2_O production^8^, we repeated the soil incubations in the laboratory, with 100 μM NO_2_^-^, using both gamma-irradiated and Zn-treated peat soil. We then deduced a sitespecific correction factor for estimates of *in-situ* rates. All our reported abiotic N_2_O production rates are therefore corrected for Zn-induced N_2_O production.

**Fig. 1.**
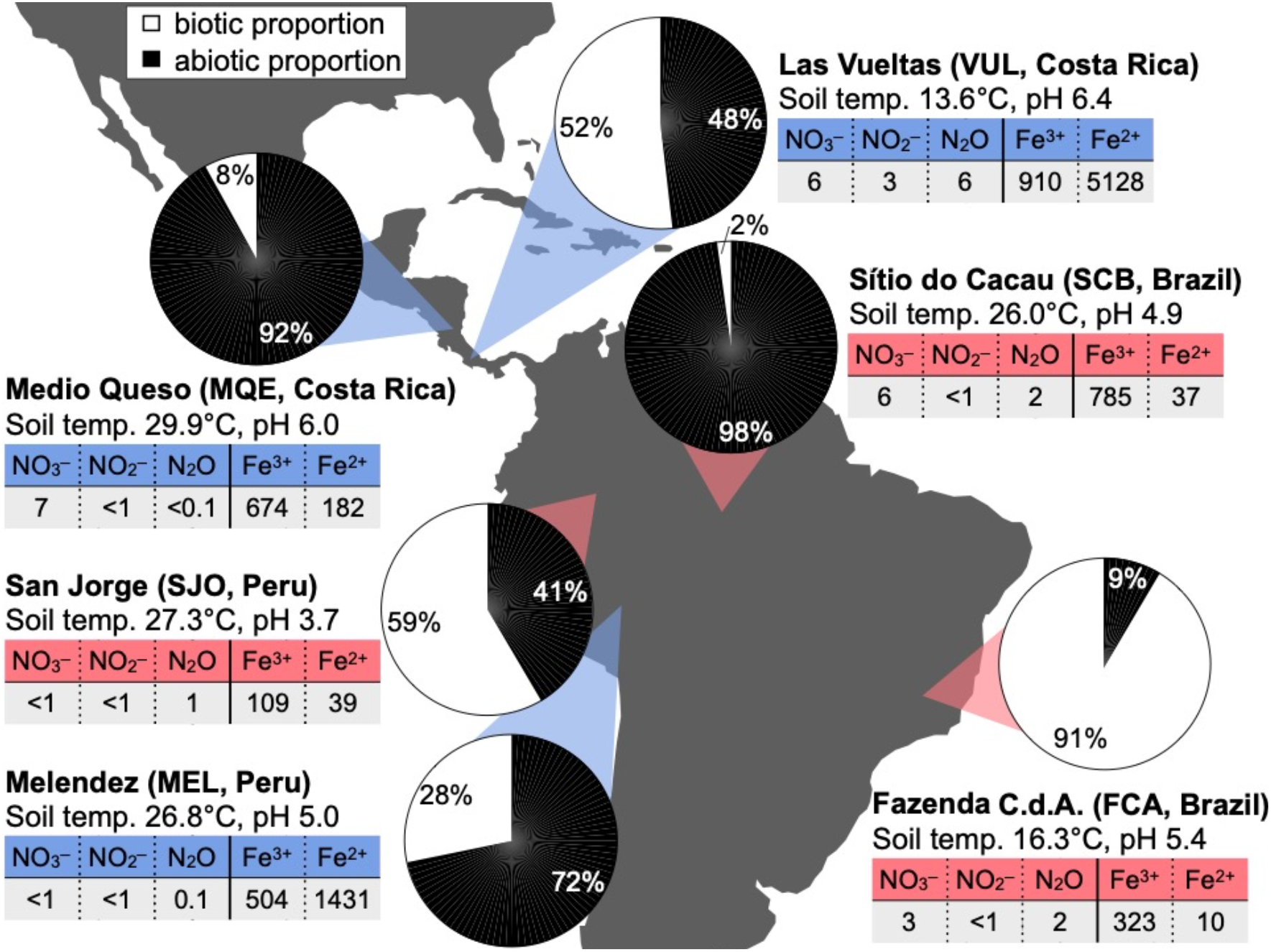
Contribution of abiotic and biotic reactions to overall N_2_O production in tropical peatlands. Rates were derived *in situ* from the enrichment of ^15^N in N_2_O after addition of ^15^NO_2_^-^ to soil in the field (n = 4). Dissolved nitrogen (measured *in-situ)* and Fe species (extracted) concentrations are given in μM. Sites are color-coded based on their NO to N_2_O yield (Table 1), showing high-NO yield (red shades) or high-N_2_O yield (blue shades).

Abiotic N_2_O production was observed in all peatlands. *In-situ* rates ranged from low (0.05-0.3 nmol N_2_O g^−1^ d^−1^) at FCA and SCB, moderate (2.4-3.3 nmol N_2_O g^−1^ d^−1^) at MQE and SJO, and high (9.2-39.0 nmol N_2_O g^−1^ d^−1^) at MEL and VUL. Abiotic N_2_O production contributed to the overall N_2_O flux to a greater extent than biotic N_2_O production at half the field sites (Fig. 1). Soil Fe^2+^ concentrations measured after extraction positively correlated with abiotic N_2_O production rates (*R^2^* = 0.99, n = 6, Supplementary Fig. 1). To determine the nitrogen yield of the chemodenitrification reaction, we incubated gamma-irradiated peat soil under anoxic conditions with 100 μM NO_2_^-^, and quantified NO_2_^-^, NO and N_2_O in time (Supplementary Fig. 2). In two peatlands (SJO, SCB), complete denitrification was achieved almost based purely on abiotic reactions (Table 1). The transformation of NO_2_^-^ into NO and N_2_O resulted in dominant yields of either product across sites, suggesting unequal nitrogen diversion directed by local peat chemistry. Our analytical approach could not confirm N_2_ as a byproduct^28^, which was presumably dominant at circum-neutral pH sites (MEL and VUL). We used this observed divergent NO_2_^-^ conversion to group the diverse peat soils into high-N_2_O (MQE, VUL, MEL) and high-NO (FCA, SJO, SCB) abiotic-yield sites (Table 1).

**Table 1.** Abiotic nitrogen yield fractions based on sterilized batch incubations. Gamma-irradiated peat soil was used for anoxic incubations initiated with the addition of 100 μM NO_2_^-^. Yield was calculated after all NO_2_^-^ was consumed and based on stable NO and N_2_O concentrations in at least two consecutive measurements. Sites with replicated samples (n = 3) included: Las Vueltas (VUL), Medio Queso (MQE), Fazenda Córrego da Areia (FCA), Melendez (MEL), Sítio do Cacau (SCB), San Jorge (SJO).

|  | SCB | SJO | FCA | MEL | VUL | MQE |
| --- | --- | --- | --- | --- | --- | --- |
| Yield in NO (%) | 97.2 | 92.8 | 74.3 | 0.2 | 0.2 | 1.3 |
| Yield in $\text{N}_2\text{O}$ (%) | 4.8 | 4.8 | 4.0 | 12.1 | 24.6 | 55.9 |
| Total yield (%) | 102 | 97.6 | 78.3 | 12.3 | 24.8 | 57.2 |

To our knowledge, this study represents the first assessment of the relative contribution of abiotic N_2_O production to the overall N_2_O production at near-natural NO_2_^-^ levels. Based on our results, abiotic reactions outcompete biotic reactions in three peatlands and are highly competitive as a source of N_2_O at another two. The measured N_2_O production rates were comparable to reported rates from a coniferous forest and grasslands^29^, although the amount of added NO_2_^-^ was at least one order of magnitude lower in our study. Relative to other evaluated ecosystems^10^, peat soils have less oxidized Fe or Mn minerals and are enriched in recalcitrant organic carbon, which would hold additional reducing power, particularly in the structurally disparate OM. For instance, pi-electron bonds are an integral part of the chemical structures found in recalcitrant organic carbon, such as phenolic or humic substances, and they are prone to interact with NO_2_^-^ (refs. ^30,31^). Besides serving as reactants, humic substances can act as regenerable electron shuttles for redox reactions in soils and sediments^32^. Iron reduction and dissolution are greatly enhanced in the presence of humic substances^33,34^, which increases the availability of Fe^2+^. The distinct production of NO and N_2_O across a gradient of Fe^2+^ concentrations suggests divergent reaction mechanisms in high-NO and high-N_2_O soils. Previous reports agree with our data that indicate the larger production of NO as the final product of chemodenitrification, which is stimulated by the self-decomposition of nitrous acid in increasingly acidic soil milieu^35,36^. High-N_2_O soils coincided with high soil Fe^2+^ abundances, and high-NO sites were associated with low Fe^2+^ (Fig. 1, Table 1). Besides Fe^2+^, the mixture of functional groups in peat OM may also be crucial in determining the NO to N_2_O balance. Except for dimethyl glyoxime and quinone oximes, oxime groups preferentially produce N_2_O and aromatics tend to produce NO^37^.

Nitrite incorporation into OM may explain lower nitrogen yields in either NO or N_2_O, in addition to the possible production of N_2_. For example, nitrosophenol (a phenol moiety that binds one NO_2_^-^ ion) tautomerizes to quinone monoxime and requires another NO_2_ ion to eject hyponitrous acid which decomposes to N_2_O^30,31^. Without the consecutive incorporation of 2 NO_2_^-^ ions, nitrosophenol remains stable, and a significant amount of NO_2_^-^ could reside in nitrosated functional groups.

### Active microbial N_2_O reduction in acidic peat soils

Concomitant to N_2_O production, we measured N_2_O consumption. Only non-sterilized samples exhibited active consumption. In sterilized samples, N_2_O was a stable end product after NO_2_^-^ addition, and ^15^N_2_ was not produced in (^15^N)_2_O amendments. Incubations with (^15^N)_2_O in the field resulted in an accumulation of the ^15^N label in N_2_ (Fig. 2) and were used to derive N_2_O reduction rates. Enrichment of ^15^N_2_ decreased with soil pH (Fig. 2), while N_2_O reduction was surprisingly observed in soils with pH as low as 3.7 (SJO). This finding is significant because N_2_O reductase assembly is post-transcriptionally inhibited by acidic pH^38^, and exposure to pH < 4 disrupts a histidine amino acid ligand to the Cu cofactor in N_2_O reductase, possibly inactivating the catalytic function (pers. com. W. Nitschke). The measured N_2_O reduction rates were higher than previously observed rates at similar acidic pH values^39^ and would extend the known physiological limits for microbial N_2_O consumption. Thus, these results demonstrate the presence and activity of N_2_O-reducing communities adapted to a wide range of peat soil pH.

**Fig. 2.**
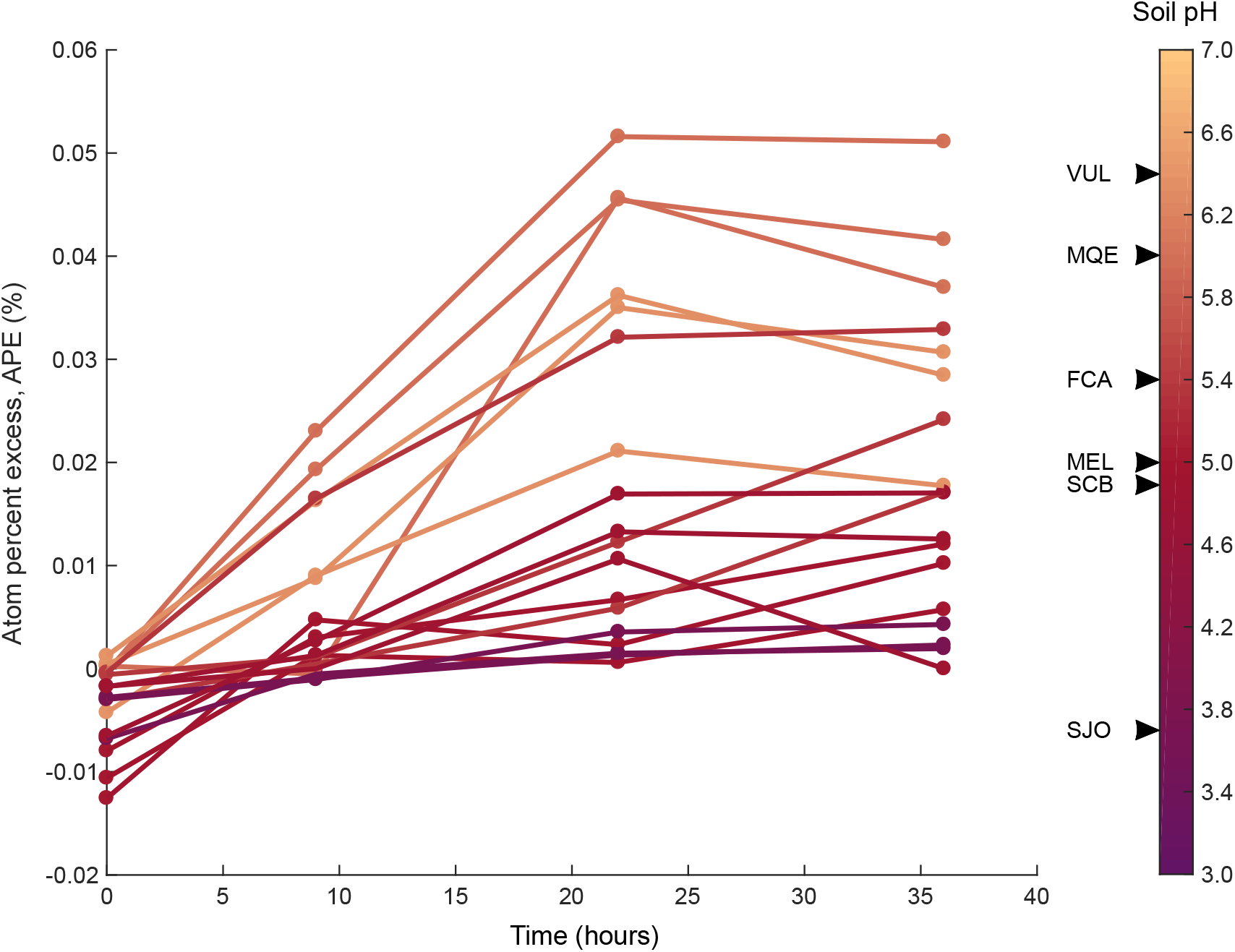
Isotopic enrichment in molecular nitrogen during *in-situ* incubations of (^15^N)_2_O with anoxic peat soil. Replicates per sites (n = 3), as listed, are colored in a gradient according to their pH. VUL, Las Vueltas; MQE, Medio Queso; FCA, Fazenda Córrego da Areia; MEL, Melendez; SCB, Sítio do Cacau; SJO, San Jorge.

### Diverse Clade II N_2_O-reducers are associated with higher N_2_O sink potential

To evaluate the relationship between the abiotic formation and microbial consumption of N_2_O, we compared reaction rates against *nosZ* gene abundances. Both processes revealed similar trends ranging from low (0.1-0.3 nmol N_2_O g^−1^ d^−1^) in SCB and FCA to moderate (0.7-1.5 nmol N_2_O g^−1^ d^−1^) in MQE and SJO to high (3-9.5 nmol N_2_O g^−1^ d^−1^) in VUL and MEL. Consumption never outpaced production, except at two sites (FCA and MEL, Fig. 3). While the variation in *nosZ* gene copies from both clades showed no significant differences among high-NO sites (FCA, SJO, SCB), they differed (ANOVA, *p* = 0.05) among high-N_2_O sites (MQE, MEL). Consumption rates gradually increased with clade II *nosZ* gene abundance at high-N_2_O sites. A clear dominance of clade II *nosZ* genes over those from clade I coincided with the elevated rates of N_2_O consumption in MEL peatland. Thus, N_2_O reducers from clade II establish an increased microbial N_2_O sink in peatlands with high abiotic N_2_O fluxes.

**Fig. 3.**
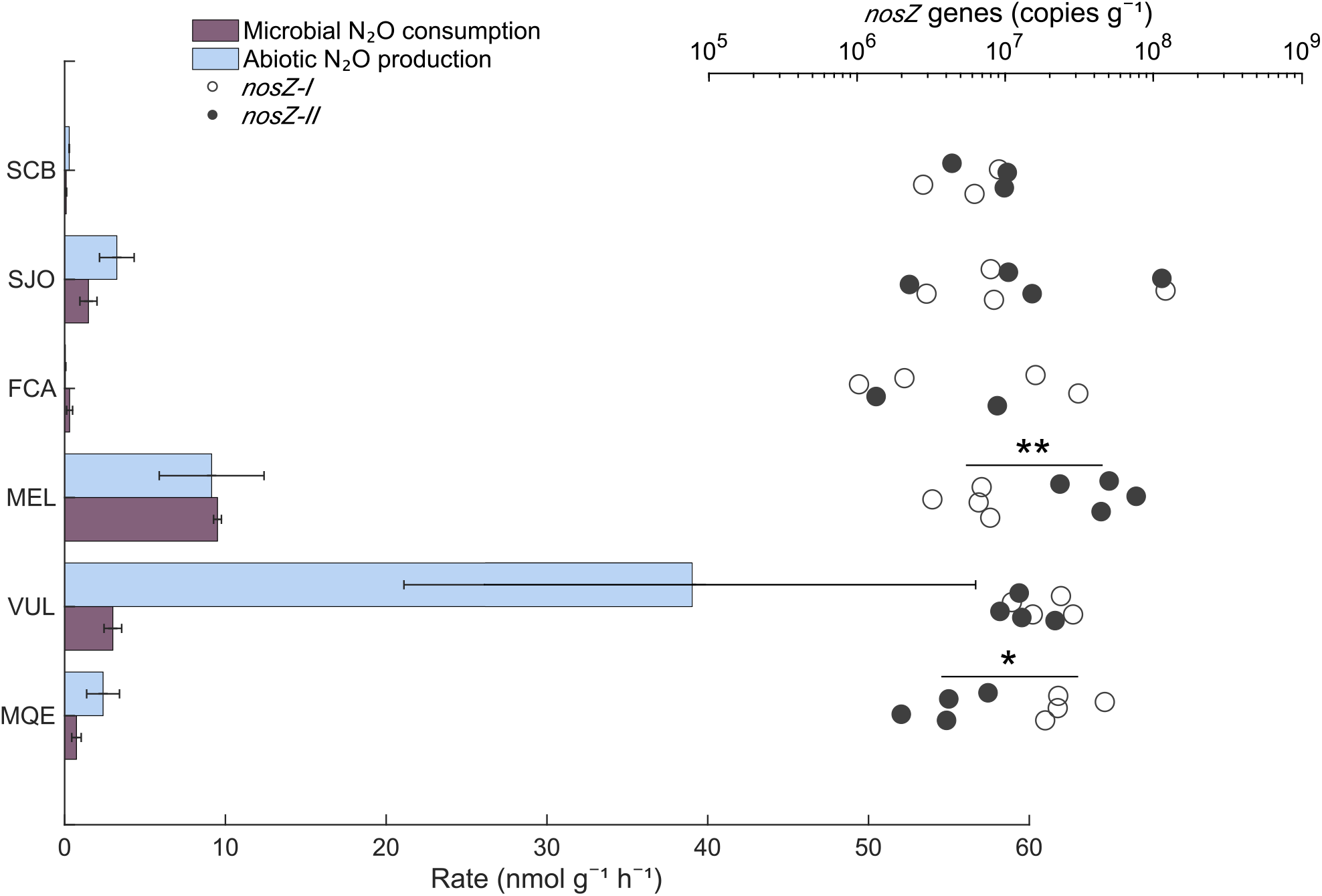
Microbial N_2_O consumption and abiotic N_2_O production (bars) along with *nosZ* gene quantities (open and filled circles). Error bars denote SD (consumption rates, n = 3; production rates, n = 4). Clade I and II *nosZ* was quantified by qPCR assays and are significantly different within sites as denoted by asterisks (ANOVA, \**p* = 0.012, \*\**p* = 0.008, n = 4). Two outliers for *nosZ-II* (SCB, ~6,200 copies g^−1^; FCA, ~590) are not shown and another two datapoints are missing due to non-amplification. See previous figures for site acronyms.

Next, to evaluate the N_2_O-reducing microbial community, we analyzed 183,265 and 80,050 taxonomically assigned *nosZ* gene amplicon sequence variants (ASVs) for clade I and II, respectively. Our analysis focused on the most abundant taxa that made up at least 1% of the total ASVs in at least one site (Fig. 4). The most abundant ASV was affiliated to the alpha-proteobacterium *Nitrospirillum amazonense* (Fig. 4). This phylotype constituted 59-64 % of clade I ASVs in the Amazon bogs SCB and SJO but was least represented in the MEL peatland (~10 %). In MEL, 23 % of ASVs belonged to *Methylocystis* species, which were also abundant in FCA (27 %). The clade II N_2_O reducers were more diverse (comprised more phyla) across all soils (Fig. 4), with *Magnetospirillum* (consistently > 10 % in high-NO sites and 30 % in VUL) and unclassified *Myxococcales* (8-50 %) as the most abundant phylotypes. To examine the observed trend of N_2_O reduction rates corresponding with *nosZ* clade II gene frequencies in the high-N_2_O sites (MEL > VUL > MQE, Fig. 3), we derived diversity indices and performed a principal component analysis (PCA). The Shannon diversity index showed a coinciding order of diversity levels (1.73 > 1.49 > 1.39, Supplementary Tables 1a-b) for high-N_2_O sites. This was also supported by a relatively high average Bray-Curtis dissimilarity of clade II *nosZ* gene sequences in MEL (0.71, Supplementary Tables 1c-d), identifying the clade II N_2_O reducer community in this peatland as most dissimilar to all others. In addition, the community structure variation among clade II was most parsimoniously explained by N_2_O consumption rates in the PCA (Supplementary Figs. 3, 4). Rather than single dominant taxa, it appeared to be the contribution of a diverse group of clade II N_2_O reducers to be responsible for the high N_2_O sink potential observed.

**Fig. 4.**
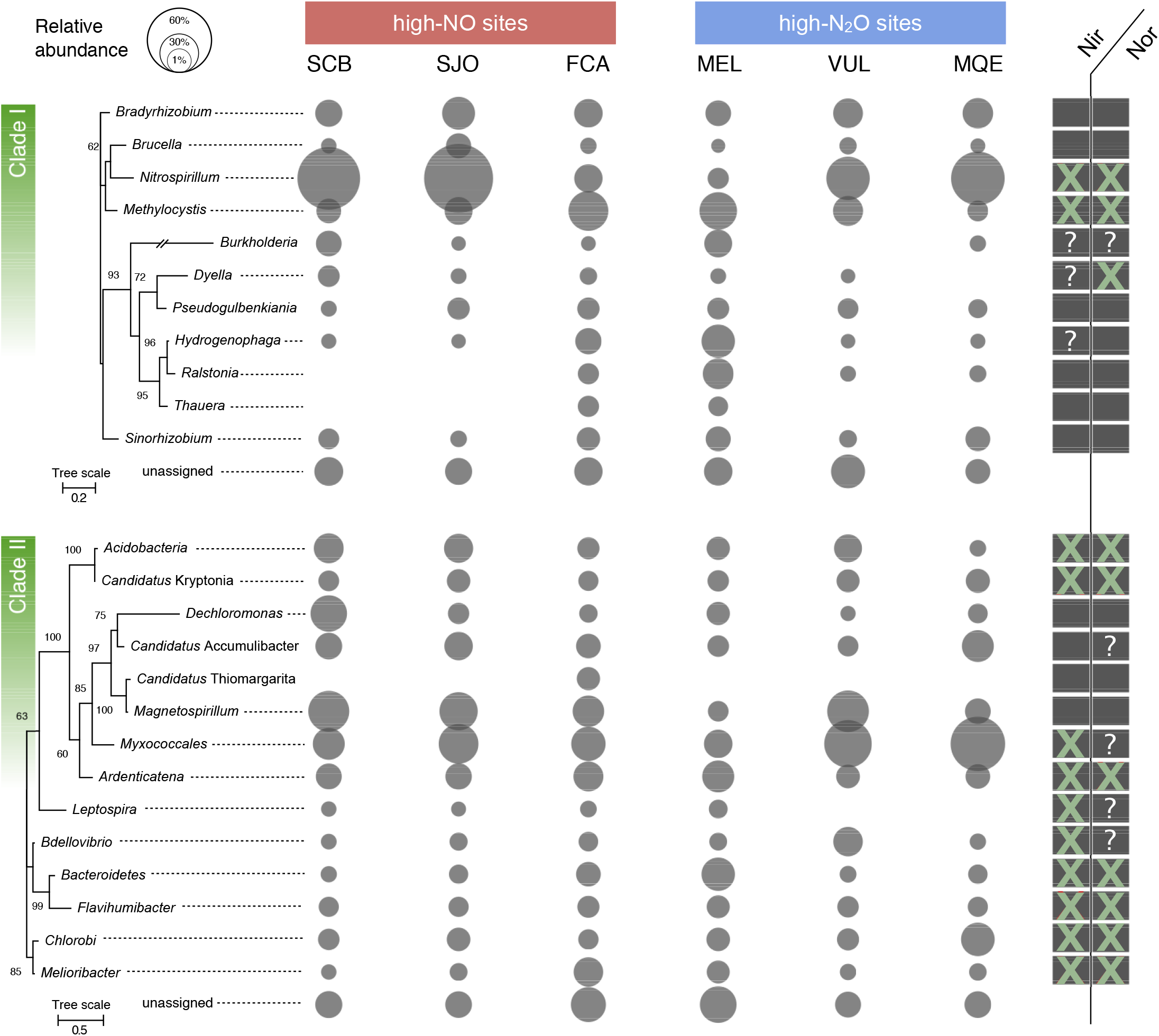
NosZ phylogeny and taxonomy in tropical peat soils. Only the most abundant amplicon sequence variants (ASVs) > 1 % were included in the analysis. Maximum likelihood phylogenetic trees are based on 1000 iterations. Nodes with 60 % or higher bootstrap support are labeled. The right panel indicates the presence (filled box) or absence (crossed box) of Nir or Nor enzyme sequences in reference proteomes. Boxes with question marks indicate an ambiguous distribution of Nir or Nor within the taxonomic group (Supplementary Table 2).

The intrinsic capacity of the peat to reduce NO_2_^-^ abiotically could provide non-denitrifying microbes that do not possess denitrification enzymes other than N_2_O reductase, called “chemodenitrifiers”^40^, an advantage over canonical denitrifiers. Chemodenitrifiers do not have to compete with chemodenitrification and simply harvest the end product N_2_O to oxidize organic substrates. Figure 4 illustrates the NO_2_^-^ reductase (either NirS or NirK) and NO reductase (NorB) enzyme repertoires present in available reference proteomes of relatives of the predicted taxa in both clades. At least 2 out of the 11 clade I taxa (including the abundant *Nitrospirillum)* and 10 out of the 14 clade II taxa indicated the absence of Nir enzymes (Fig. 4). Half of the clade II reference proteomes were missing both Nir and Nor proteins. Importantly, the Myxococcales ASVs showed no differences in abundance among high-NO sites but gradually increased, similar to the N_2_O yield, in the high-N_2_O sites. This order, which also includes *Anaeromyxobacter dehalogenans,* the hallmark organism of clade II N_2_O reducers^22,41^, is frequently represented in acidic, organic-rich soils^40,42^, presumably with abiotic NO and N_2_O production potential. Further, other bacteria such as *Dechloromonas^4^,, Ardenticatena^44^,* and *Melioribacter^4^* also mediate iron reduction, an additional trait that could promote chemodenitrification by recycling Fe^2+^. Therefore, chemodenitrifiers may outcompete canonical denitrifiers in the studied peatlands (e.g., *Nitrospirillum)* and abundance patterns of the Myxococcales suggest a notable benefit for some chemodenitrifiers in soils associated with high abiotic N_2_O yields.

Another abundant taxon of the N_2_O-reducing community was *Magnetospirillum* (clade II) that includes several iron-oxidizing species. These alpha-proteobacteria synthesize the mixed-valence iron mineral magnetite, which can accumulate in soils, also under the influence of abiotic crystallization^46^. Secondary iron mineral formation can be widespread in the tropics, driven by dissolved and particulate iron originating from weathering and desilication^47–49^. Ferrous iron-bearing minerals, such as magnetite, can serve as catalysts for NO_2_^-^ reduction by providing reaction sites at the mineral surface^50,51^. The possession of an N_2_O reductase makes sense for *Magnetospirillum,* assuming cells are associated with, or at least grow in proximity to, magnetite. Analyses of the iron phases present would be necessary to follow up on this in more detail, which was off the scope of our study. We also acknowledge that our insight into the temporal activity response is limited. Information on the actual *in-situ* transcription levels is needed to better assess how the clades react to fluctuating abiotic pulses of N_2_O^52^.

## Discussion

### Decoupling of abiotic N_2_O production and microbial N_2_O consumption

Our data show the concomitant occurrence of both abiotic N_2_O production and microbial consumption (Fig. 3 and Supplementary Fig. 2), and their positive correlation points to the coupling of both processes in several sites (Supplementary Fig. 5). However, the mountain bog site (VUL) exhibited unusually high abiotic production rates and relatively low consumption rates (Fig. 3). A lower soil temperature than in the other peatlands and the differential sensitivity of production and consumption (Supplementary Table 3 and Supplementary Text) could lead to such kinetic effects^53–55^. Along these lines, the decoupling of production and consumption establishes the potential for vast N_2_O emissions when changing environmental conditions impart selectively negative effects on consumption. For instance, while peatland drainage occurs naturally between wet and dry seasons^56^, N_2_O cycling could become decoupled by aerobic conditions created by extended peatland drainage with microbial denitrification persisting only in anoxic microsites. Nitrite, fueled by increased nitrification, could still be abiotically reduced to N_2_O because acidic peat soil stabilizes Fe^2+^ via two mechanisms. First, Fe^2+^ oxidation by oxygen is kinetically hindered at low pH. Oxidation rates are significantly slowed at pH ≤ 6.5 ^57^, a pH regime applying to most peatlands, including all in our study. Second, Fe^2+^ complexed by OM is resilient to oxidation. Experimental evidence suggests that tannic acid^58^, phenolics^59^, or natural humic acid^60^ stabilize the Fe^2+^ pool in the presence of oxygen by the formation of a redox-buffering shell^60^ and re-reduction of Fe^3+^. However, little is known concerning how Fe^2+^ complexation affects the NO_2_^-^ accessibility and reduction. Nevertheless, these previous findings indicate that the reactants for chemodenitrification are sufficiently available even at higher oxygen concentrations (> 6 μM), leading to a potential predominance of abiotic N_2_O production over biotic N_2_O production.

We present evidence that active abiotic-biotic N_2_O cycling is prevalent in tropical peatlands, where denitrification is not a purely biological pathway, but rather a “mosaic” of biotic and abiotic reduction reactions (Fig. 5). Furthermore, our results support the idea that functional modularity complements not only the interrelationship of microbial groups but also concomitant interactions between microbes and spontaneous chemical reactions. Abiotic N_2_O formation in tropical peatlands can have important regional consequences in the context of observed N_2_O fluxes and higher rates in response to drainage^61^, and putatively drought^62^, as well as possible effects on reducing organic carbon release^63^. For example, compared to the other soils, abiotic N_2_O production was moderate in SJO, an acidic oligotrophic site, showing a net production of 1.8 nmol N_2_O g^−1^ day^−1^. With the estimation of the global extent of acidic oligotrophic tropical peatlands alike SJO at 1,003,719 km^2^ (ref. ^15^), this could amount to a total depth-integrated abiotic N_2_O flux ranging from 0.1 to 3.9 Tg N_2_O yr^−1^ depending on the depth of NO_2_^-^ diffusion. This contribution is a major uncertainty that could account for two percent to more than half of all combined terrestrial N_2_O fluxes from natural sources of South America, Africa, and Southeast Asia. With a factor of 298 g CO_2_ -equivalents per g N_2_O over a 100 year time^64^, abiotic N_2_O fluxes could also alter the net radiative effect of tropical peatlands. Accordingly, by bypassing heterotrophic respiration through chemodenitrification, less organic carbon is mineralized to CO_2_. Considering ~50 mg CO_2_-C m^-2^ yr^−1^ emitted from SJO-like tropical peatlands^7^ and 4 moles of N_2_O required to mineralize 1 mole of organic carbon, chemodenitrification would suppress at least 2.7 % of the organic carbon mineralization, promoting the retention of 2 Tg C yr^−1^ across oligotrophic tropical peatlands. These estimates are conservative because they do not include processes such as diversion of nitrogen oxides into NO, nitrosation of OM, and the consumption deficit observed in the high-altitude peatland. Sensitivity to lower temperatures could also impede microbial N_2_O reduction in northern peatlands, which would imply an imbalanced cycling of N_2_O and substantial N_2_O release from abiotic origins.

**Fig. 5.**
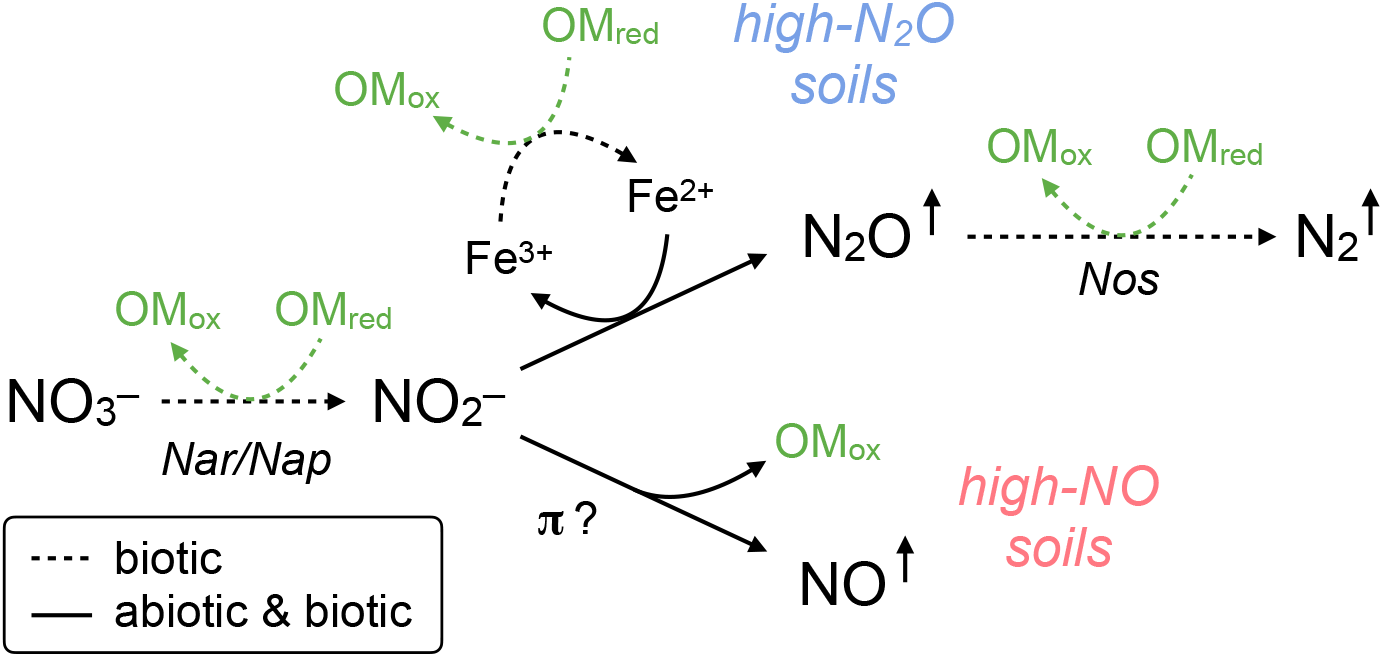
Schematic representation of denitrification pathways in tropical peatlands. Nitrate (NO_3_^-^) reduction to NO_2_^-^ occurs at significant rates only with catalysis by NO_3_^-^ reductases. Nitrite is rapidly reduced via abiotic and biotic reactions. At lower pH (≤ 5.4), NO is the dominant product. Nitrosation into OM may be an alternative abiotic process in soils with minor nitrogenous gas production reliant on organic compounds containing pi-electron bonds (π). In Fe^2+^-rich peat, N_2_O is the dominant product, involving Fe redox cycling that can fuel dissimilatory Fe reduction^11^. The only N_2_O consumption pathway in peat soil is N_2_O reductase-dependent reduction to N_2_, which is active even in the most acidic soils tested (pH 3.7). All related heterotrophic reactions induce oxidation of OM (OMred → OMox) and eventually peat carbon mineralization.

## Methods

### Study sites

Six peatlands were chosen to cover a geochemical spectrum, including acidic (pH 3.7-5) soils, low (10 μM) and high (> 5 mM) Fe^2+^ concentrations, varying OM content and soil temperature (Supplementary Table 4). Most sites were under little to no anthropogenic influence (Supplementary Table 5), with two exceptions: Fazenda Córrego da Areia (FCA) located within a catchment experiencing agricultural run-off in Brazil, and Medio Queso (MQE) in a Costa Rican river delta surrounded by agricultural run-off and cattle raising. The San Jorge (SJO) peatland is located in the Pastaza-Marañón foreland basin and Melendez (MEL) is in the Madre de Dios river terraces, both in Peru. Sítio do Cacau (SCB) is located in Central Amazonia (Amanã Reserve) in Brazil. Las Vueltas (VUL), located in Costa Rica’s cloud forests of the Cerro Las Vueltas Reserve, differed most drastically from the other sites due to its higher altitude (2,500 m a.s.l.). Field work was conducted in September 2017 (Peru) and between April (Costa Rica) and July (Brazil) in 2018.

### ^15^N tracer experiment in the field

Colorimetric assays to determine ambient soil NO_2_^-^ and NO_3_^-^ concentrations were performed in the field using a YSI 9500 portable spectrophotometer (YSI Inc.), including reagent kits, according to the manufacturer’s instructions. Dissolved N_2_O was sampled by collecting pore water into a pre-evacuated vial and subsequent degassing by shaking for 5 minutes. Thereafter, headspace was transferred into a pre-evacuated vial and stored underwater prior to analysis with a GC-ECD. Soil temperature and pH were measured with a YSI A10 pH probe (Ecosense, YSI).

Anaerobic glove bags filled with argon (Ar) were used to provide anaerobic conditions in the field while distributing soil into glass incubation vials (160 mL). Topsoils (10 cm) were sampled with 30 ml-barrel customized plastic corers. Inside the glove bag, the center 5 cm (~15 g) soil was diluted 1:5 (w/v) into vials with anoxic water extracted directly from the deep peat soil via a water line connected to the glove bag. Separate sample sets received ^15^NO_2_^-^ (label fraction = 0.1, Cambridge Isotopes) at 10× the soil ambient NO_2_^-^ concentration and doubly labeled (^15^N)_2_O (label fraction = 1.0, Cambridge Isotopes) at 5× the soil ambient N_2_O concentration. Thus, the amount of ^15^N tracer applied varied slightly between sites but reflected a similar order of magnitude. ^15^NO_2_^-^ incubations included untreated and 87.5 mM zinc chloride-poisoned (ZnCl_2_, Fisher Scientific) soils in replicates of four (n = 4). Soil slurries were incubated in insulating containers to avoid temperature fluctuations, and gas samples were taken for (^15^N)_2_O analysis at the beginning of incubation and after 24 h (n = 4), and for ^30^N_2_ analysis at four time points spread over 36 h (n = 3). Gas sampling was destructive (entire headspace used) for (^15^N)_2_O analysis or by replacement with 5 mL Ar gas for ^30^N_2_ analysis. The sample times for the ^30^N_2_ analysis were adapted from a previous study^65^. We also prepared zinc-poisoned (^15^N)_2_O incubations to test for abiotic N_2_O consumption. The gas samples were stored underwater in borosilicate glass vials closed with thick butyl rubber stoppers^66^ prior to analysis at Michigan State University. Isotopic compositions of N_2_O and N_2_ were measured using an Elementar Isoprime isotope ratio mass spectrometer (IR-MS) interfaced with an Elementar TraceGas chromatographic system. Rate calculations closely followed a previously developed and tested protocol^14^. Given the constraints of sterilant applications in the field, we repeated the zinc-amended incubations in the lab, complementarily to incubations with gamma-irradiated soils. The rates from both experiments were used to calculate a correction factor accounting for artifacts caused by the zinc addition^8^. The rates derived in the field were then multiplied by the correction factor (Supplementary Fig. 6 and Supplementary Table 6). The Brazilian sites SCB and FCA have associated data from gamma-irradiated soil, but data from zinc-treated soil are missing because of logistic issues concerning the shipment of non-sterilized (not gamma-irradiated) soil. The final rates were combined according to the following equation for net *insitu* N_2_O formation:

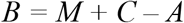

where *B* is the biotic N_2_O production rate, *M* is the mixed rate (from untreated ^15^NO_2_^-^ incubations), *C* is the microbial N_2_O consumption rate [from (^15^N)_2_O incubations], and *A* is the abiotic N_2_O production rate (from poisoned ^15^NO_2_^-^ incubations).

### Laboratory incubations

In an anoxic glove box (0.1% H_2_ for O_2_ reduction in N_2_), gamma-sterilized and zinc-treated soil were prepared separately. Gamma sterilization followed a previous method^8^. Zinc chloride was applied, as described for field experiments. Roots and coarse particles (> 5 mm in diameter) were removed from soils, and soils were slurried 1:5 (w/v) in anoxic, sterile 18.2 MΩocm water. The slurry was homogenized before equal quantities were distributed into culture vials and sealed with sterile butyl rubber stoppers. An anoxic and filter-sterilized NO_2_^-^ solution was injected (final concentration 100 μM) into microcosms that were previously flushed with pure N_2_. The microcosms were agitated briefly to disperse the added substrate and then kept under dark, static conditions at room temperature for ~100 h. Dissolved NO_2_^-^ was quantified with the Griess reagent (Promega, Kit G2930), and NO and N_2_O were analyzed as described below.

### Soil Fe measurements

Soils for Fe analysis were kept in anoxic serum bottles and refrigerated during transport. The species Fe^2+^ and Fe^3+^ were extracted and separated as previously described^8^ and quantified in acidified aqueous solution by inductively coupled plasma-optical emission spectrometry (ICP-OES; Thermo iCAP6300 at the Goldwater Environmental Laboratory at Arizona State University). The ICP-OES pump rate for the Ar carrier was set to 50 rpm, and Fe2395 and Fe2599 lines were used for Fe quantification. Iron concentrations were determined from a calibration curve (0.01-10 mg L^−1^) by diluting a standard solution (100 mg L^−1^, VHG Labs, product no. SM75B-500) in 0.02 N HNO_3_.

### N_2_O gas measurements

Using a gas-tight syringe (VICI Precision Sampling), 200 μL of gas sample was injected into a gas chromatograph (GC, SRI Instruments) equipped with an electron-capture detector (ECD). Two continuous HayeSep-D columns were kept at 90 °C (oven temperature), and N_2_ (UHP grade 99.999 %, Praxair Inc.) was used as carrier gas. The ECD current was 250 mV, and the ECD cell was kept at 350 °C. The N_2_O measurements were calibrated over a range of 0.25-100 ppmv using customized standard mixtures (Scott Specialty Gases, accuracy ±5 %). Gas concentrations were corrected for solubility effects using Henry’s law and the dimensionless concentration constant *k_H_^cc^(N_2_O*) = 0.6112 to account for gas partitioning into the aqueous phase at 25 °C and 1 atm^67^.

### NO gas measurements

Nitric oxide (NO) was quantified in the microcosm headspace with a chemiluminescence-based analyzer (LMA-3D NO_2_ analyzer, Unisearch Associates Inc., Concord, Canada). Headspace gas (50 μL) was withdrawn with a CO_2_-N_2_-flushed gas-tight syringe and injected into the analyzer. The injection port was customized to fit the injection volume and consisted of a T-junction with an air filter at one end and a septum at the other end. An internal pump generated consistent airflow. Our method followed a previous protocol^68^, with minor adjustments. Briefly, NO was oxidized to NO_2_ by a CrO_3_ catalyst. The NO_2_ was carried across a fabric wick saturated with a Luminol solution (Drummond Technology Inc., Canada). Readings were corrected for background NO_2_ every 15 min (“zeroing”). Shell airflow rate was kept at 500 mL min^−1^, and the span potentiometer was set to 8. Measurements were calibrated with a 0.1 ppm NO (in N_2_) standard (< 0.0005 ppm NO_2_, Scott-Marin, Riverside, CA, USA) over a range of 5-1,000 ppbv. Gas concentrations were corrected using Henry’s law and the dimensionless concentration constant *k_H_^cc^*(NO) = 0.0465 to account for gas partitioning into the aqueous phase at 25 °C and 1 atm^67^.

### Molecular analyses

Peat samples from four randomly distributed locations (coinciding with incubation locations) within a peatland were collected and frozen at −20 °C for subsequent DNA extraction. Genomic DNA was extracted using a NucleoSpin Soil DNA extraction kit (Macherey-Nagel GmbH, Düren, Germany).

For quantitative polymerase chain reactions (qPCR), we used two primer pairs and a total reaction volume of 15 μL with 1.5 μL DNA template (35-50 ng genomic DNA). The clade I *nosZ* gene was amplified with PowerUp SYBR Green master mix (Applied Biosystems, Foster City, CA), to which 3 mM MgCl2 was added. Forward and reverse primer concentrations were 1 μM, and previous cycler conditions were used^69^. The clade II *nosZ* gene was amplified using SYBR Fast, ROX low master mix (Kapa Biosystems, Roche Holding AG, Basel, Switzerland), and 1.2 μM primer concentration^70^. Thermal cycling was initiated with 3 min of denaturing at 95 °C, followed by 40 cycles of the following stages: 30 s at 95 °C, 60 s at 58 °C, 30 s at 72 °C, 30 s at 80 °C, and a final melting-curve. Samples were run in technical duplicates on 96-well plates using a Quantstudio 3 thermocycler (Applied Biosystems, Foster City, CA). Standards were prepared using linearized plasmids. Standard curves indicated efficiencies of 94 % *(R^2^* = 0.99, *nosZ* clade I) and 85 % (*R*^2^ = 0.99, *nosZ* clade II) and melting curves showed no detectable primer dimers over the duration of 40 amplification cycles.

For Illumina amplicon sequence analysis, we developed independent *nosZ* clade I and II libraries. PCR-amplification of both *nosZ* genes used the Promega GoTaq qPCR kit (Promega, Madison, WI) and 1 μL of DNA template (25-50 ng genomic DNA) in a total reaction volume of 20 μL. Targeting the clade I *nosZ* gene, we used a novel primer pair^71^. The reaction mix included 0.2 mg mL^−1^ bovine serum albumin (BSA) and 0.8 μM primer concentration. For the clade II *nosZ* gene, we used the same primer as used for qPCR in reactions of 1 mg mL^−1^ BSA and 0.8 μM primer concentration. Cycling conditions for clade II *nosZ* amplification were used as described^70^. Thermal cycling conditions for clade I *nosZ* amplification were an initial 2 min denaturing step at 95 °C, followed by 33 cycles of 95 °C for 45 s, annealing by 53 °C for 45 s, and a 72 °C extension for 30 s, and a final extension at 72 °C for 7 min. Amplification was verified by gel electrophoresis using 1 % agarose in 1 Tris-acetate-EDTA buffer. Samples were multiplexed^72^, normalized (SequalPrep kit #1051001, Invitrogen), and submitted for sequencing to the DNASU core facility at Arizona State University, with 2× 300-bp paired-end Illumina MiSeq.

Paired-end sequences were merged and demultiplexed, then we used the USEARCH pipeline^73^ to 1) correct strand orientations, 2) sort out singletons, and 3) denoise the dataset. We used *alpha* = 2 for a stringent denoising of sequences^74^ because reads were not clustered with any identity radius to obtain ASVs. The sequences were translated and frameshift-corrected by Framebot^75^ with low sequence loss (< 10 %). The amino acid sequences obtained were classified using Diamond^76^ version 0.9.25. The search was conducted in Diamond’s *sensitive* mode, with an e-value cutoff of 10^-5^, resulting in the top 5 % hits. Sequences were parsed through two databases; the NCBI database RefSeq (release 95) containing 146,381,777 non-redundant protein sequences and manually curated databases built from 2,817 (clade I) and 2,929 (clade II) sequences off the FunGene repository^77^ using the search parameters 80 % HMM coverage and a minimum length of 550 amino acids. The taxonomy achieved with the curated databases was used for downstream analysis because of a higher number of classified sequences. The output was imported into Megan^78^ version 6.18.0, where a weighted lowest common ancestor (LCA) algorithm [default parameters according to Huson et al.^79^] was run to assign a single taxonomic lineage to each read. ASV tables were pasted into Krona^80^ for visual inspection of results. Reads with abundances of > 1 % in at least one site were extracted, and consensus sequences were determined for each taxonomic group. Maximum-likelihood phylogenetic trees were constructed with consensus sequences in Mega X^81^. To infer presence/absence of Nir and Nor enzymes in representative proteomes, UniProt reference (manually curated) proteomes were screened using BlastP with default parameters. NirS (Q51700, *Paracoccus denitrificans* PD1222), NirK (O31380, *Bradyrhizobium japonicum),* NorB (Q51663, *Paracoccus denitrificans)* were used as amino acid query sequences.

The *nosZ* sequences have been deposited in the GenBank, EMBL, and DDBJ databases as SRA Bioproject XXX.

### Statistical analyses

All statistical tests were performed with JMP Pro software (Version 13.1.0, SAS Institute Inc.). Analysis of variance (ANOVA) was used with *p* < 0.05 to test significantly different values for gene quantities across soils. Plotting and regression analysis were done with the Matlab R2018a software package (Version 9.4.0.813654, Mathworks Inc.).

## Data availability

All data to evaluate the conclusions of the study are present in the paper and the Supplementary Information.

## Acknowledgments

We acknowledge Dr. Rodil Tello Espinoza, Dr. Tedi Pacheco Gomez, David Reyna, Brian Crnobrna, Dr. Outi Lähteenoja, Kelsen Arbaiza, Antenor Hurtado Carmona, Paulo Fonteboa, Ronaldo César Chaves, Juan Rodrigo Trucios, Cely Mariela Cadillo-Quiroz, the UFSJ Graduate Program in Geography (PPGEOG) and Office for International Affairs (ASSIN/UFSJ) for assistance and help during stages of field work. We also thank Wolfgang Nitschke (CNRS/BIP) for discussions and Mohamed Abdalla for his work supporting this effort at the USAID-GDR program at ASU.

This study was primarily funded by an NSF-DEB award (#1355066) and a SOLS-KED ASU award (ECR A548 HC) to H.C-Q, a Global Development Research Scholarship to S.B. and H.C-Q in partnership with the USAID-Global Development Lab and Peruvian and Brazilian USAID missions. S.B. also received support by the Lewis & Clark Fund for Exploration and Field Research in Astrobiology provided by the American Philosophical Society (APS). N.E.O. was funded in part by the DOE Great Lakes Bioenergy Research Center (DOE BER Office of Science DE-SC0018409).

## Author contributions

S.B., N.E.O., and H.C.-Q. designed the study. S.B. conducted the field work with essential contributions from A.G.P-C, G.P.P., J.D.U.-M., L.P.R., J.M.F.M., I.G.B., and B.G.. S.B., M.F.O., A.F.S., and M.C. R. performed laboratory experiments and molecular analyses. S.J.H. supported the NO analysis. K.E.H. conducted soil gamma sterilization. C.R.P. supported qPCR analysis. H.G. analyzed isotopic abundances of gas samples. S.B., I.G.B., B.G., N.E.O., and H.C.-Q. performed the data analysis. S.B. and H.C.-Q. wrote the manuscript, and all co-authors contributed to the final version of the paper.

