## Supplementary Information for "Coupled abiotic-biotic cycling of nitrous oxide in tropical peatlands"

**This PDF file includes:**

Supplementary Text
Supplementary Figures 1 to 6
Supplementary Tables 1 to 6
References

### Supplementary Text

*Comparing activation energies involved in abiotic-biotic $N_2O$ cycling*

Supplementary Table 3 illustrates the divergence between activation energies i) of abiotic and biotic/mixed $N_2O$ production reactions and ii) of abiotic $N_2O$ production and consumption. Abiotic $N_2O$ production reactions are highly competitive against biotic/mixed reactions because of low activation energies (Supplementary Table 3) of the abiotic process. In contrast, abiotic consumption warrants more than ten times the amount of activation energy needed to biotically produce $N_2O$ (per mole), making enzymatic $N_2O$ consumption crucial to determine the $N_2O$ sink in soils. Because enzymatic activity is more sensitive to temperature changes[1], $N_2O$ consumption might be more affected by temperature than $N_2O$ production. We explain the offset of abiotic $N_2O$ production and biotic $N_2O$ consumption in the mountain bog VUL (Fig. 3) by slowed microbial $N_2O$ reduction rates at lower temperatures due to altitude. The cause of this effect could be translated to high-latitude sites. In the case of global warming, rising temperatures may close the production/consumption offset and decrease $N_2O$ emissions but increase organic carbon respiration and $CO_2$ emissions. Our interpretation is therefore in line with macro-level models based on metabolic theory. These models predict ecosystem organic carbon respiration to increase proportionately more than primary production because of higher activation energies of respiratory processes[2,3].

**Supplementary Figures**

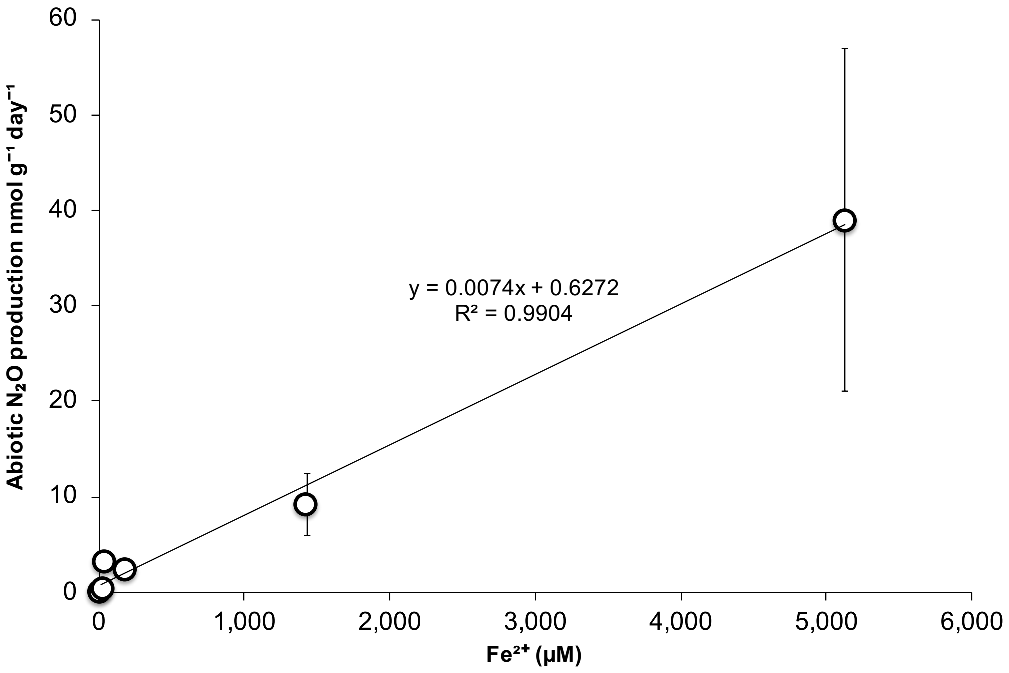


**Supplementary Fig. 1.** Relationship between extracted $Fe^{2+}$ concentrations and *in-situ* abiotic $N_2O$ production rates (n = 4).

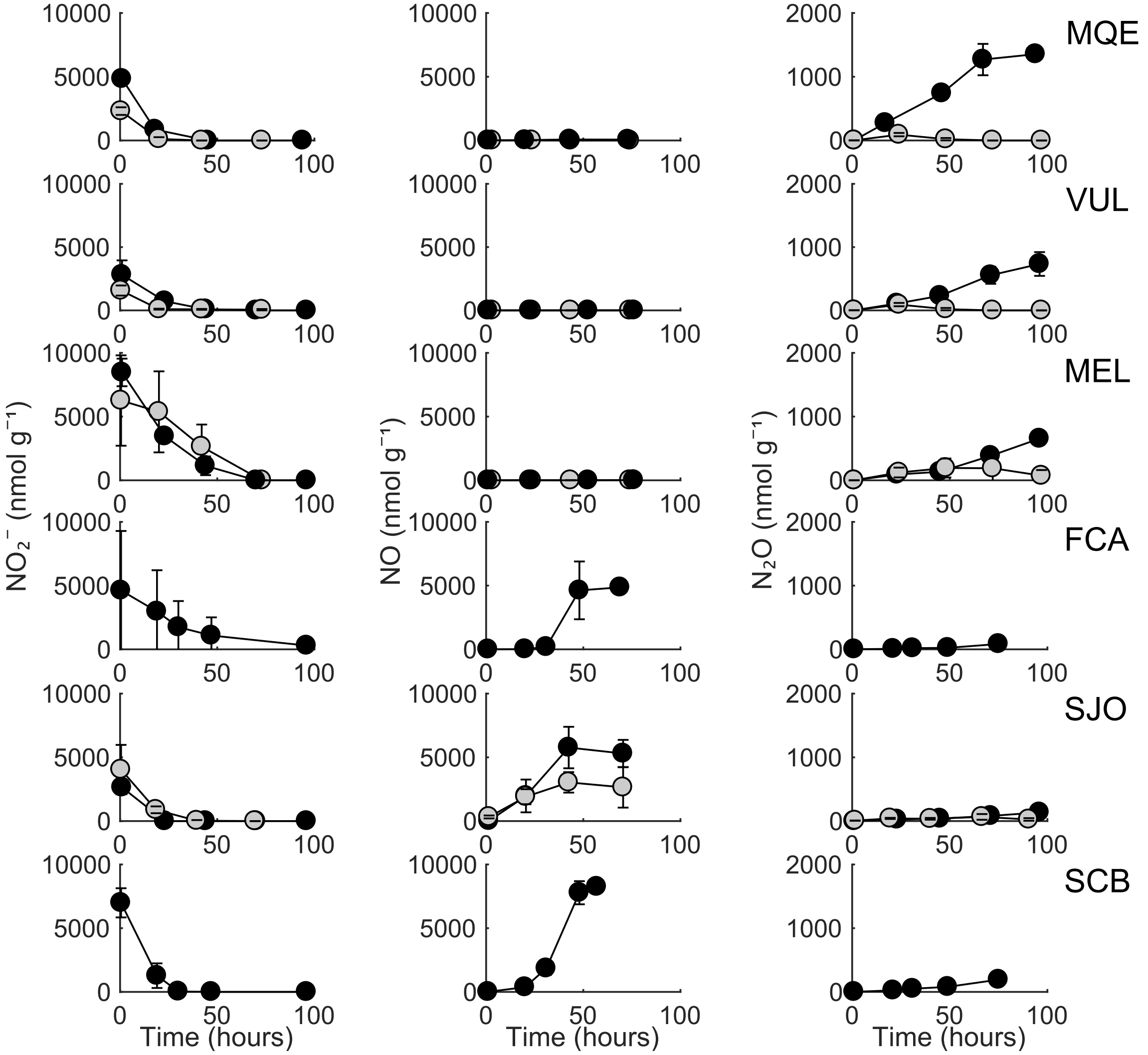


**Supplementary Fig. 2. Chemodenitrification assays.** Liquid phase and headspace from gamma-irradiated (black) and untreated (gray) batch incubations were sampled for analysis of $NO_2^-$, NO and $N_2O$ after initial amendment with 100 µM $NO_2^-$. Yield was calculated using the concentration average of the last two measurements. Units were normalized to soil dry weight. Error bars denote *SD* (n = 4).

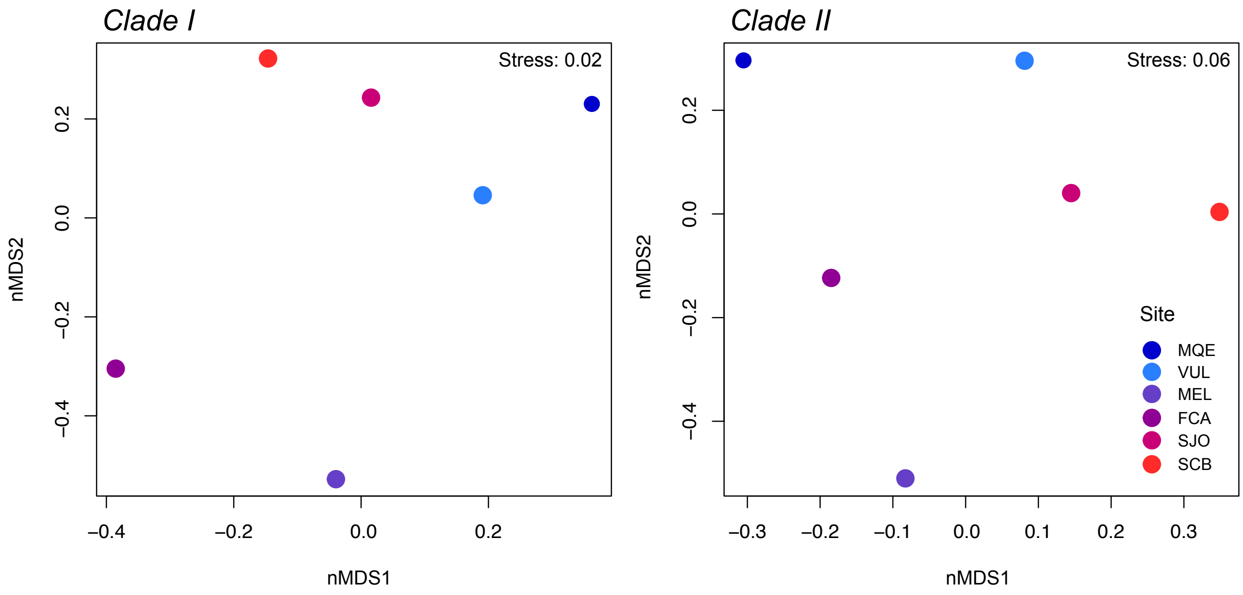


**Supplementary Fig. 3. Non-metric multidimensional scaling (nMDS) plot of the taxonomic composition of clade I and II $N_2O$ reducers in the six study sites.** Bray-Curtis similarity distances are calculated from pairwise taxonomic profile comparisons between sites (Tables S1c-d). Data points are based on 4 replicate samples at each site.

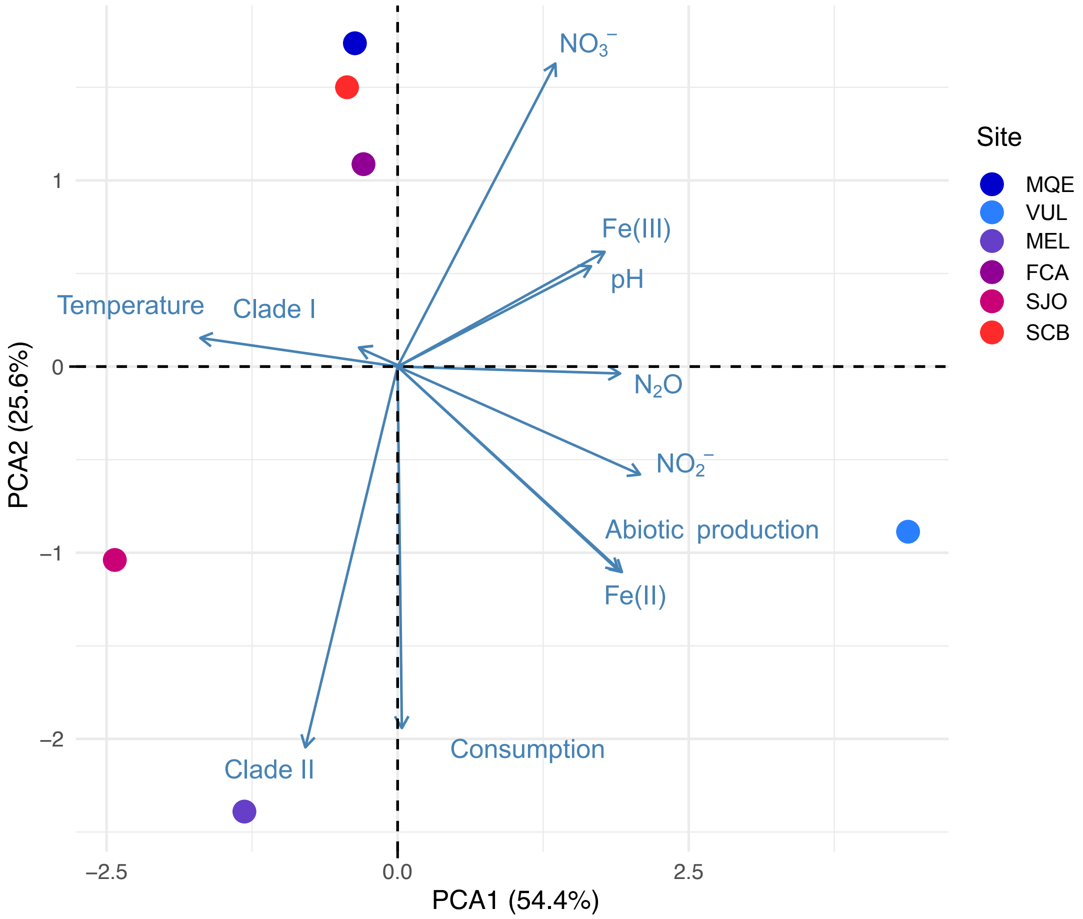


**Supplementary Fig. 4. Principle component analysis (PCA) correlation biplot showing the scores of the six study sites (filled circles), and the factor loadings of environmental variables (arrows).** The first two principal components of the PCA account for 80.0% of the total variation among sites. *Clade I, Clade II* = Clade I and II gene frequencies determined by qPCR; *Abiotic production, consumption* = measured $N_2O$ abiotic production and $N_2O$ consumption rates; all chemical compounds refer to their concentration measurements across sites. Data were $\log_{10}(x+1)$-transformed, centered, and scaled. Data points are based on 4 replicate samples at each site.

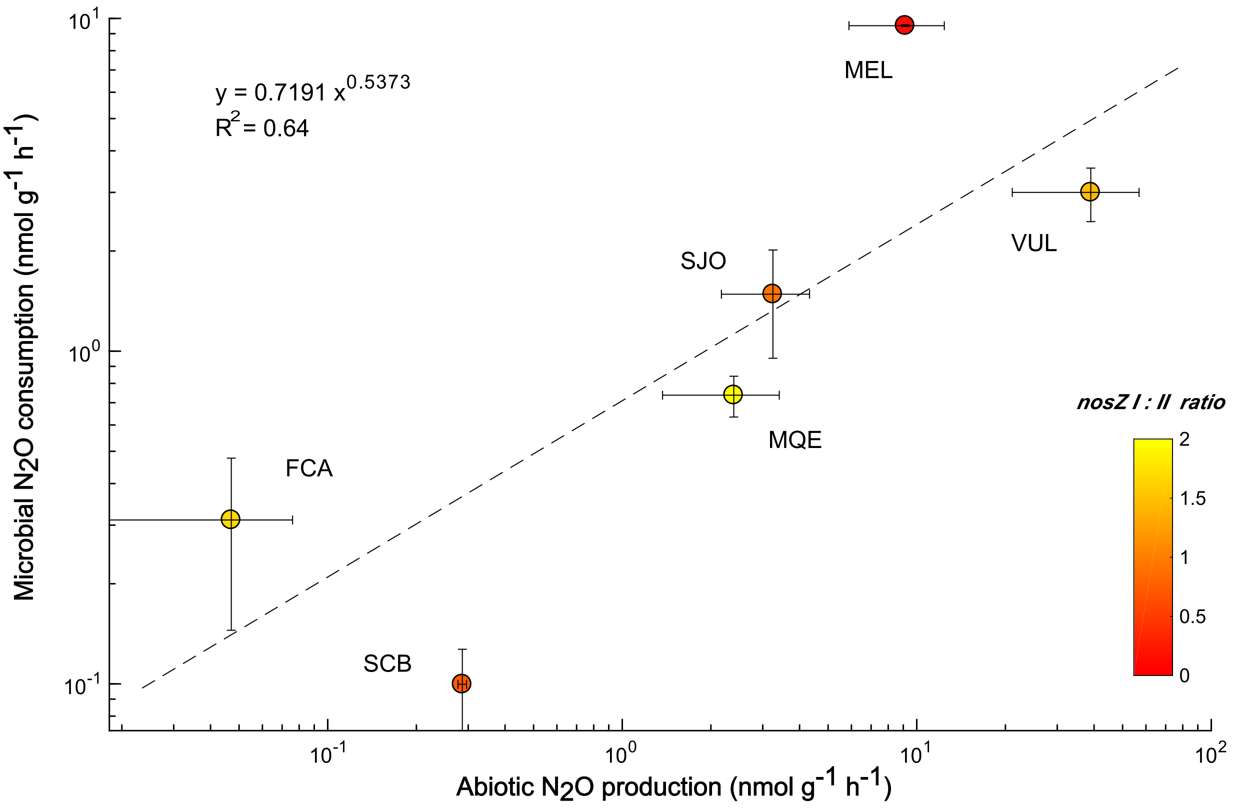


**Supplementary Fig. 5. Relation of abiotic $N_2O$ production (x-axis) and microbial $N_2O$ consumption (y-axis).** *NosZ* clade I:II ratios < 2 are marked with a color gradient. We infer a coupling of $N_2O$ production and consumption based on the positive correlation shown here. *NosZ* clade I:II ratios > 1 (FCA, MQE, VUL) are associated with lower $N_2O$ consumption rates as abiotic production increases (shallow slope). In turn, *nosZ* clade I:II ratios < 1 (SCB, SJO, MEL) are associated with higher $N_2O$ consumption rates as abiotic production increases (steep slope), indicating increased $N_2O$ sink potential due to higher abundance of clade II $N_2O$ reducers.

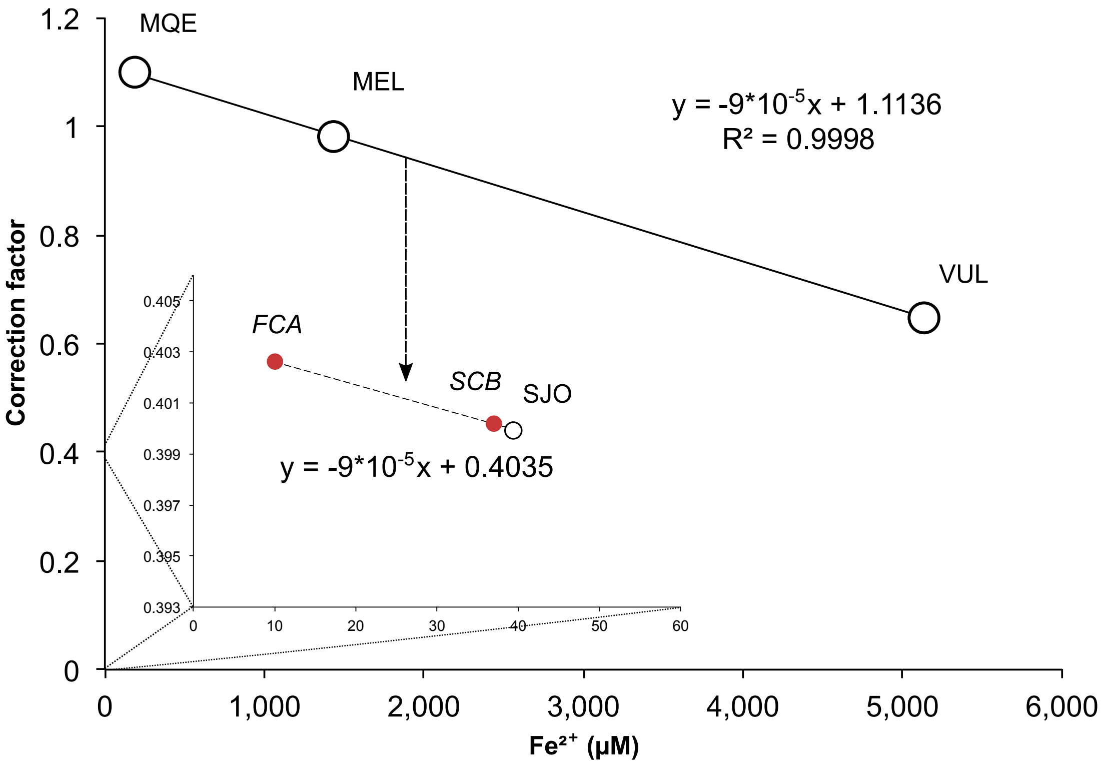


**Supplementary Fig. 6. Derivation of the correction factors for sites FCA and SCB.** The correction factor (y) was derived by constructing a line with slope $-9*10^{-5}$ through the data point SJO and using $Fe^{2+}$ concentrations from FCA and SCB. For this, we assumed a linear relationship between $Fe^{2+}$ concentration and $N_2O$ production rates within the high-NO and high-$N_2O$ group, respectively. The correction factor values are similar within the high-NO group due to relatively small differences in $Fe^{2+}$ and a shallow slope of the regression line.

### Supplementary Tables

**Supplementary Table 1a.** Alpha diversity indices of *nosZ* amplicon sequences affiliated to clade I.

| Site | Reads | Berger-Parker | Buzas-Gibson | Chao-1 | Dominance | Equitability | Jost | Robbins | Simpson | Shannon* |
|---|---|---|---|---|---|---|---|---|---|---|
| *SJO* | 13035 | 0.79 | 0.000173 | 9 | 0.364 | 0.371 | 1.77 | 0 | 0.636 | 0.816 |
| *MEL* | 17849 | 0.707 | 0.000143 | 9 | 0.467 | 0.427 | 2.11 | 0.2 | 0.533 | 0.939 |
| *SCB* | 7256 | 0.51 | 0.000633 | 11 | 0.687 | 0.636 | 3.71 | 0.0833 | 0.313 | 1.52 |
| *FCA* | 24561 | 0.341 | 0.000234 | 11 | 0.785 | 0.729 | 5.07 | 0.0833 | 0.215 | 1.75 |
| *VUL* | 29426 | 0.617 | 9.36E-05 | 9 | 0.553 | 0.461 | 2.43 | 0 | 0.447 | 1.01 |
| *MQE* | 36927 | 0.759 | 5.82E-05 | 9 | 0.393 | 0.348 | 1.81 | 0 | 0.607 | 0.765 |

**calculated using log to base e.*

**Supplementary Table 1b.** Alpha diversity indices of *nosZ* amplicon sequences affiliated to clade II.

| Site | Reads | Berger-Parker | Buzas-Gibson | Chao-1 | Dominance | Equitability | Jost | Robbins | Simpson | Shannon* |
|---|---|---|---|---|---|---|---|---|---|---|
| *SJO* | 10899 | 0.339 | 0.000515 | 13 | 0.781 | 0.673 | 4.96 | 0 | 0.219 | 1.73 |
| *MEL* | 8329 | 0.338 | 0.000675 | 13 | 0.766 | 0.673 | 4.75 | 0.0714 | 0.234 | 1.73 |
| *SCB* | 14512 | 0.276 | 0.000514 | 14 | 0.829 | 0.761 | 6.43 | 0 | 0.171 | 2.01 |
| *FCA* | 5008 | 0.308 | 0.0014 | 13 | 0.804 | 0.759 | 5.79 | 0 | 0.196 | 1.95 |
| *VUL* | 14569 | 0.424 | 0.000305 | 12 | 0.701 | 0.601 | 3.71 | 0 | 0.299 | 1.49 |
| *MQE* | 15296 | 0.549 | 0.000262 | 12 | 0.643 | 0.559 | 3.22 | 0 | 0.357 | 1.39 |

**calculated using log to base e.*

**Supplementary Table 1c.** Bray-Curtis dissimilarity matrix based on *nosZ* amplicon sequences affiliated to clade I.

| | SCB | SJO | VUL | MQE | FCA | MEL |
|---|---|---|---|---|---|---|
| SCB | 0 | | | | | |
| SJO | 0.227 | 0 | | | | |
| VUL | 0.435 | 0.273 | 0 | | | |
| MQE | 0.519 | 0.41 | 0.272 | 0 | | |
| FCA | 0.724 | 0.708 | 0.668 | 0.859 | 0 | |
| MEL | 0.847 | 0.831 | 0.707 | 0.87 | 0.586 | 0 |

**Supplementary Table 1d.** Bray-Curtis dissimilarity matrix based on *nosZ* amplicon sequences affiliated to clade II.

| | SCB | SJO | FCA | MEL | VUL | MQE |
|---|---|---|---|---|---|---|
| SCB | 0 | | | | | |
| SJO | 0.297 | 0 | | | | |
| FCA | 0.484 | 0.346 | 0 | | | |
| VUL | 0.428 | 0.36 | 0.426 | 0 | | |
| MQE | 0.69 | 0.527 | 0.556 | 0.458 | 0 | |
| MEL | 0.71 | 0.656 | 0.593 | 0.786 | 0.789 | 0 |

**Supplementary Table 2a. BlastP results for reference proteomes belonging to clade I.** Percentage identities (% ID) are provided for each alignment with NirS, NirK, and NorB query sequences (Methods). Low-identity (< 50 %) hits were additionally checked for their annotation in the proteome and using associated primary literature. Values in brackets indicate hits that could not be confirmed (concluding absence). If all BlastP alignments within a taxonomic group suggested either absence or presence, the results were consistent. If the results were ambiguous, it was not possible to conclude absence/presence of the enzyme.

| ASV classification (Fig. 4) | Select UniProt reference proteome | Organism ID | NirS (% ID) | NirK (% ID) | NorB (% ID) | Presence/absence in ref. proteomes |
|---|---|---|---|---|---|---|
| Bradyrhizobium | *Bradyrhizobium diazoefficiens* JCM 10833 | 224911 | n.d. | 99.45 | 63.07 | consistent |
| | *Bradyrhizobium* sp. LMTR 3 | 189873 | n.d. | 74.73 | 63.6 | |
| | *Bradyrhizobium* sp. STM 3843 | 551947 | n.d. | 90.83 | 61.47 | |
| | *Bradyrhizobium* sp. ORS 278 | 114615 | n.d. | 87.6 | 62.34 | |
| | *Bradyrhizobium* sp. Leo170 | 1571199 | n.d. | 74.59 | 63.66 | |
| | *Bradyrhizobium erythrophlei* GAS478 | 1437360 | (31.63) | 88.64 | 24.38 | |
| | *Bradyrhizobium elkanii* UASWS1015 | 29448 | n.d. | 74.03 | 62.12 | |
| Brucella | *Brucella abortus* strain 2308 | 359391 | n.d. | 71.73 | 65.46 | consistent |
| | *Brucella anthropi* ATCC 49188 | 439375 | n.d. | 71.43 | 64.5 | |
| | *Brucella suis* bv. 1 (strain 60) | 644346 | n.d. | 71.43 | 64.56 | |
| | *Brucella thiophenivorans* DSM 7216 | 571255 | n.d. | 71.43 | 64.29 | |
| Nitrospirillum | *Nitrospirillum amazonense* CBAmc | 1441467 | n.d. | n.d. | n.d. | consistent |
| | *Nitrospirillum amazonense* Y2 | 1003237 | n.d. | n.d. | n.d. | |
| Methylocystis | *Methylocystis* sp. SC2 | 187303 | n.d. | n.d. | (24.14) | consistent |
| | *Methylocystis bryophila* S285 | 655015 | n.d. | n.d. | n.d. | |
| | *Methylocystis heyeri* H2 | 391905 | n.d. | n.d. | (25.65) | |
| Burkholderia | *Burkholderia multivorans* ATCC 17616 | 395019 | n.d. | n.d. | n.d. | ambiguous |
| | *Burkholderia pseudomallei* K96243 | 272560 | n.d. | 34.26 | 27.35 | |
| | *Burkholderia* sp. SRS-W-2-2016 | 1926878 | n.d. | n.d. | n.d. | |
| | *Burkholderia* sp. JS23 | 1770053 | n.d. | n.d. | n.d. | |
| | *Burkholderia* sp. Leaf177 | 1736287 | (30.77) | n.d. | n.d. | |
| | *Burkholderia glumae* BGR1 | 626418 | n.d. | n.d. | n.d. | |
| | *Candidatus* Burkholderia verschuerenii UZHbot4 | 242163 | n.d. | n.d. | n.d. | |
| | *Candidatus* Burkholderia pumila UZHbot3 | 1090375 | n.d. | n.d. | n.d. | |
| | *Burkholderia* sp. CCGE1003 | 640512 | n.d. | n.d. | n.d. | |
| | *Burkholderia* sp. DHOD12 | 2571746 | n.d. | n.d. | n.d. | |

| | | | | | | |
|---|---|---|---|---|---|---|
| Dyella | *Dyella* sp. M7H15-1 | 2501295 | n.d. | n.d. | n.d. | ambiguous (NirK) |
| | *Dyella japonica* DSM 16301 | 1440762 | (33.82) | (33.61) | n.d. | |
| | *Dyella japonica* A8 | 1217721 | n.d. | (34.08) | n.d. | |
| | *Dyella* sp. OK004 | 1855292 | n.d. | n.d. | n.d. | |
| | *Dyella soli* KACC 12747 | 522319 | n.d. | 35.25 | n.d. | |
| | *Dyella psychrodurans* 4MSK11 | 1927960 | (37.93) | n.d. | n.d. | |
| | *Dyella tabacisoli* L4-6 | 2282381 | n.d. | n.d. | n.d. | |
| | *Dyella choica* 4M-K27 | 1927959 | (28.87) | (26.61) | n.d. | |
| | *Dyella thiooxydans* ATSB10 | 445710 | n.d. | 38.08 | (26.84) | |
| Pseudogulbenkiania | *Pseudogulbenkiania ferrooxidans* 2002 | 279714 | 59.86 | (32.11) | 56.95 | consistent |
| | *Pseudogulbenkiania subflava* DSM 22618 | 1123014 | 59.35 | (35) | 57.17 | |
| | *Pseudogulbenkiania* sp. NH8B | 748280 | 64.23 | (32.11) | 57.17 | |
| Hydrogenophaga | *Hydrogenophaga* sp. T4 | 1437444 | n.d. | n.d. | 54.71 | ambiguous (NirS) |
| | *Hydrogenophaga* sp. PBC | 795665 | n.d. | n.d. | 26.82 | |
| | *Hydrogenophaga* sp. A37 | 1945864 | 60.54 | n.d. | 53.7 | |
| | *Hydrogenophaga* sp. H7 | 1882399 | 57.62 | n.d. | 54.22 | |
| | *Hydrogenophaga* sp. IBVHS1 | 1985169 | n.d. | n.d. | 25.1 | |
| | *Hydrogenophaga crassostreae* LPB0072 | 1763535 | 60.24 | n.d. | 51.68 | |
| Ralstonia | *Ralstonia* sp. A12 | 1217052 | n.d. | 34.07 | 25.85 | consistent |
| | *Ralstonia solanacearum* GMI1000 | 267608 | n.d. | 33.52 | 27.74 | |
| | *Ralstonia metallidurans* CH34 | 266264 | 64.56 | n.d. | 27.73 | |
| | *Ralstonia eutropha* ATCC 17699 | 381666 | 63.36 | n.d. | 27.22 | |
| Thauera | *Thauera terpenica* 58Eu | 1348657 | 54.03 | n.d. | 57.42 | consistent |
| | *Thauera* sp. K11 | 2005884 | 53.9 | n.d. | 100 | |
| | *Thauera* sp. 63 | 497321 | 61.43 | n.d. | 57.98 | |
| | *Thauera* sp. 27 | 305700 | 66.03 | n.d. | 56.77 | |
| | *Thauera linaloolentis* 47Lol / DSM 12138 | 1123367 | 52.87 | n.d. | 57.75 | |
| | *Thauera* sp. MZ1T | 85643 | 53.79 | n.d. | 57.08 | |
| | *Thauera chlorobenzoica* 3CB1 | 96773 | 53.6 | n.d. | 56.43 | |
| Sinorhizobium | *Sinorhizobium fredii* USDA 257 | 1185652 | n.d. | 74.1 | 64.43 | consistent |
| | *Sinorhizobium fredii* GR64 | 882101 | n.d. | 69.23 | 64.33 | |
| | *Sinorhizobium meliloti* strain 1021 | 266834 | n.d. | 69.02 | 64.21 | |

**Supplementary Table 2b. BlastP results for reference proteomes belonging to clade II.** Percentage identities (% ID) are provided for each alignment with NirS, NirK, and NorB query sequences (Methods). Low-identity (< 50 %) hits were additionally checked for their annotation in the proteome and using associated primary literature. Values in brackets indicate hits that could not be confirmed (concluding absence). If all BlastP alignments within a taxonomic group suggested either absence or presence, the results were consistent. If the results were ambiguous, it was not possible to conclude absence/presence of the enzyme.

| ASV classification (Fig. 4) | Select UniProt reference proteome | Organism ID | NirS (% ID) | NirK (% ID) | NorB (% ID) | Presence/absence in ref. proteomes |
|---|---|---|---|---|---|---|
| Acidobacteria | *Granulicella pectinivorans* DSM 21001 | 474950 | n.d. | n.d. | n.d. | consistent |
| | *Granulicella tundricola* ATCC BAA-1859 | 1198114 | n.d. | n.d. | n.d. | |
| | *Acidipila dinghuensis* DHOF10 | 1560006 | n.d. | n.d. | n.d. | |
| | *Edaphobacter dinghuensis* EB95 | 1560005 | n.d. | n.d. | n.d. | |
| | *Edaphobacter aggregans* EB153 | 570835 | n.d. | (25.45) | (25.84) | |
| Candidatus Kryptonia | *Candidatus* Kryptobacter tengchongensis JGI-24 | 1643429 | n.d. | n.d. | n.d. | consistent |
| | *Candidatus* Kryptonium thompsoni | 1633631 | n.d. | n.d. | n.d. | |
| Dechloromonas | *Dechloromonas* sp. HYN0024 | 2231055 | 53.42 | n.d. | 55.73 | consistent |
| | *Dechloromonas denitrificans* ATCC BAA-841 | 281362 | 65.17 | n.d. | 56.18 | |
| | *Dechloromonas* sp. | 1917218 | 67.56 | (33.72) | 56.99 | |
| Candidatus Accumulibacter | *Candidatus* Accumulibacter sp. SK-12 | 1454001 | 56.25 | n.d. | n.d. | ambiguous (NorB) |
| | *Candidatus* Accumulibacter sp. SK-02 | 1453999 | 61.21 | n.d. | n.d. | |
| | *Candidatus* Accumulibacter sp. BA-93 | 1454004 | 60.51 | n.d. | 26.71 | |
| | *Candidatus* Accumulibacter sp. BA-94 | 1454005 | 24.93 | n.d. | n.d. | |
| | *Candidatus* Accumulibacter sp. BA-91 | 1454002 | 61.21 | n.d. | n.d. | |
| | *Candidatus* Accumulibacter sp. SK-11 | 1454000 | 56.25 | n.d. | n.d. | |
| | *Candidatus* Accumulibacter aalborgensis | 1860102 | 64.56 | n.d. | n.d. | |
| Candidatus Thiomargarita | *Candidatus* Thiomargarita nelsonii | 1003181 | 59.08 | n.d. | 56.5 | inconclusive |
| Magnetospirillum | *Magnetospirillum aberrantis* SpK | 908842 | 58.7 | n.d. | 65.1 | consistent |
| | *Magnetospirillum magneticum* AMB-1 | 342108 | 64.24 | n.d. | 64.75 | |
| | *Magnetospirillum* sp. UT-4 | 2681467 | 57.45 | n.d. | 63.84 | |
| | *Magnetospirillum* sp. SS-4 | 2681465 | 66.18 | n.d. | 65.42 | |

| | | | | | | |
|---|---|---|---|---|---|---|
| | *Magnetospirillum moscoviense* BB-1 | 1437059 | 55.83 | n.d. | 62.61 | |
| Myxococcales | *Polyangium fumosum* DSM 14668 | 889272 | n.d. | (26.88) | (25.06) | ambiguous (NorB) |
| | *Anaeromyxobacter dehalogenans* 2CP-C | 290397 | n.d. | (31.87) | 29.62 | |
| | *Polyangium* sp. SDU3-1 | 2567896 | n.d. | n.d. | (26.37) | |
| | *Cystobacter ferrugineus* Cbfe23 | 83449 | n.d. | n.d. | (26.06) | |
| | *Sorangiineae bacterium* NIC37A_2 | 1970199 | n.d. | n.d. | n.d. | |
| | *Stigmatella aurantiaca* DW4/3-1 | 378806 | n.d. | n.d. | n.d. | |
| | *Vulgatibacter incomptus* DSM 27710 | 1391653 | n.d. | n.d. | n.d. | |
| | *Minicystis rosea* DSM 24000 | 888845 | n.d. | n.d. | n.d. | |
| Ardenticatena | *Ardenticatena maritima* strain 110S | 872965 | n.d. | (34.29) | n.d. | inconclusive |
| Leptospira | *Leptospira interrogans serovar* strain 56601 | 189518 | n.d. | (31.54) | n.d. | ambiguous (NorB) |
| | *Leptospira interrogans serovar* HAI135 | 1085538 | n.d. | (32.31) | n.d. | |
| | *Leptospira biflexa serovar* ATCC 23582 | 456481 | n.d. | (33.23) | 45.43 | |
| | *Leptospira licerasiae serovar* VAR 010 | 1049972 | n.d. | n.d. | 46.92 | |
| | *Leptospira weilii* strain Ecochallenge | 1049986 | n.d. | (32.31) | n.d. | |
| | *Leptospira borgpetersenii* strain 200701203 | 1193007 | n.d. | (31.54) | n.d. | |
| | *Leptospira ryugenii* YH101 | 1917863 | n.d. | n.d. | n.d. | |
| Bdellovibrio | *Bdellovibrio exovorus* JSS | 1184267 | (41.82) | n.d. | n.d. | ambiguous (NorB) |
| | *Bdellovibrio bacteriovorus* ATCC 15356 | 264462 | (23.57) | (31.15) | 47.69 | |
| | *Bdellovibrio bacteriovorus* W | 765869 | n.d. | (33.54) | n.d. | |
| | *Bdellovibrio bacteriovorus* R0 | 959 | n.d. | (34.22) | 47.69 | |
| | *Bdellovibrio* sp. SKB1291214 | 1732569 | (28.71) | n.d. | n.d. | |
| | *Bdellovibrio* sp. NC01 | 2220073 | n.d. | (23.37) | n.d. | |
| Bacteroidetes | Bacteroidetes bacterium UKL13-3 | 1690483 | (31.36) | n.d. | n.d. | consistent |
| | Bacteroidetes bacterium SCGC AAA795-G10 | 2175803 | (29.58) | n.d. | n.d. | |
| | Bacteroidetes bacterium B1(2017) | 2015582 | (29.59) | n.d. | (23.9) | |
| Flavihumibacter | *Flavihumibacter* sp. ZG627 | 1463156 | n.d. | n.d. | (25.65) | consistent |
| | *Flavihumibacter petaseus* NBRC 106054 | 1220578 | n.d. | (48.15) | n.d. | |
| | *Flavihumibacter solisilvae* strain 3-3 | 1349421 | (23.26) | (48.15) | (46.64) | |
| | *Flavihumibacter soli* R14 | 2705549 | n.d. | n.d. | n.d. | |

| | | | | | | |
|---|---|---|---|---|---|---|
| Chlorobi | *Chlorobium tepidum* ATCC 49652 | 194439 | n.d. | n.d. | n.d. | consistent |
| | *Chlorobium ferrooxidans* DSM 13031 | 377431 | n.d. | n.d. | n.d. | |
| | *Chlorobium phaeobacteroides* DSM 266 | 290317 | n.d. | n.d. | n.d. | |
| | *Chlorobium limicola* DSM 245 | 290315 | (30.56) | n.d. | (23.49) | |
| | *Chlorobium chlorochromatii* CaD3 | 340177 | n.d. | n.d. | n.d. | |
| | *Chlorobium phaeovibrioides* DSM 265 | 290318 | n.d. | n.d. | n.d. | |
| | *Pelodictyon phaeoclathratiforme* DSM 5477 | 324925 | n.d. | n.d. | n.d. | |
| Melioribacter | *Melioribacter* sp. GWF2_38_21 | 1801612 | n.d. | n.d. | n.d. | consistent |
| | *Melioribacter roseus* JCM 17771 | 1191523 | n.d. | n.d. | n.d. | |

**Supplementary Table 3.** Activation energies of abiotic, mixed, or biotic $N_2O$ production/consumption reactions.

| Process | Mechanism | Activation energy (kJ $mol^{-1}$) | Reference |
|---|---|---|---|
| $N_2O$ production | abiotic | 18.4 | 4 |
| | mixed | 133.2 | 5 |
| | mixed | 102.2 | 5 |
| | mixed | 57.6 | 5 |
| | mixed | 84 | 6 |
| $N_2O$ consumption | abiotic | 250 | 7 |
| | mixed | 28 | 8 |
| | mixed | 49 | 8 |
| | mixed | 57 | 8 |
| | mixed | 59 | 8 |
| | mixed | 60 | 8 |
| | biotic | 54.4 | 9 |
| | biotic | 10.5 | 10 |

**Supplementary Table 4.** Soil geochemistry and select elemental concentrations (n = 3).

| | **Medio Queso (MQE)** | **Las Vueltas (VUL)** | **Melendez (MEL)** | **Fazenda C.d.A (FCA)** | **San Jorge (SJO)** | **Sítio Cacau (SCB)** |
|---|---|---|---|---|---|---|
| Soil temp. (°C) | 29.85±2.1 | 13.6±0.1 | 26.8±0.7 | 16.3±0.1 | 27.3±0.4 | 26±0.2 |
| pH | 6.0±0.1 | 6.4±0.005 | 5.0±0.01 | 5.3±0.1 | 3.7±0.03 | 4.9±0.1 |
| OM fraction (%) | 42.0±1.4 | 76.8±0.5 | 24.9±1.5 | 79.4±16 | 24.2±0.7 | 21.1±1.5 |
| Al (ppm) | 166±13 | 1362±38 | 321.45±25 | 295.2±18 | 362.6±25 | 179±15 |
| Ca (ppm) | 1047.5±49 | 755.4±59 | 961.7±38 | 795.2±35 | 994.4±64 | 634.2±23 |
| Co (ppm) | 0.5±0.5 | 0.7±0.7 | 0.9±0.9 | 0.6±0.6 | 0.4±0.4 | 1±1 |
| Cr (ppm) | 13.2±0.6 | 6.1±2.2 | 6.6±0.9 | 5±0.4 | 7.5±0.4 | 4.3±0.5 |
| Cu (ppm) | 4±0.7 | 11.2±0.5 | 7.7±1.9 | 12.8±2.7 | 7±0.6 | 4±2.5 |
| K (ppm) | 340.9±48 | 336.5±1.6 | 1193.3±522 | b.d. | 463.7±412 | 423.7±160 |
| Mg (ppm) | 645.5±57 | 749.4±51 | 1353.5±61 | 673.3±339 | 722.1±4.3 | 199.1±94 |
| Mn (ppm) | 21.8±0.3 | 234.3±1.8 | 62.1±0.7 | 6.3±0 | 8.7±0.1 | 7.4±0.1 |
| Mo (ppm) | 0.2±0.2 | 6±0.7 | 1.2±1.2 | 0.9±0.9 | 0.9±0.9 | 2.1±1.6 |
| Na (ppm) | 103.4±4 | 175.2±91 | 1903.5±20 | b.d. | 1361±67 | b.d. |
| Ni (ppm) | 7.6±0.5 | 4.7±0.6 | 4.4±1.1 | 3.4±0.8 | 4.5±0.7 | 3.6±1.2 |
| Sr (ppm) | 3.6±1 | 2.5±0.3 | 4.4±0.7 | 1.4 | 25.6±2 | 2.3±0.6 |
| V (ppm) | 7.8±6.1 | 14±5.4 | 9±6.9 | 19.7±7.4 | 9.2±7.5 | 9.1±7.6 |
| Zn (ppm) | 8131.5±12 | 4250±15 | 4441±25 | 5217±24 | 4456±11 | 3207±18 |

**Supplementary Table 5. Study site coordinates.**

| Site | Code | Latitude (°) | Longitude (°) |
|---|---|---|---|
| San Jorge, Iquitos, Peru | SJO | -4.058 | -73.189 |
| Melendez, Puerto Maldonado, Peru | MEL | -12.467 | -69.178 |
| Sítio do Cacau, Tefé, Brazil | SCB | -2.626 | -64.593 |
| Fazenda Córrego da Areia, Prados, Brazil | FCA | -21.024 | -44.086 |
| Las Vueltas, Dota, Costa Rica | VUL | 9.624 | -83.848 |
| Medio Queso, Los Chiles, Costa Rica | MQE | 11.038 | -84.687 |

**Supplementary Table 6. Correction of *in-situ* $N_2O$ production rates.** To relate $N_2O$ production rates from zinc-treated to gamma-irradiated soil, two sets of incubations were conducted under controlled conditions in the laboratory and production rates were measured. The correction factor was then applied to *in-situ* production rates from zinc-treated soil.

| *Site* | Rates in zinc-treated incubations (nmol $N_2O$ $g^{-1}$ $h^{-1}$) | Rates in gamma-irradiated incubations (nmol $N_2O$ $g^{-1}$ $h^{-1}$) | Correction factor |
|---|---|---|---|
| *MQE* | 14.1 | 15.5 | 1.1 |
| *VUL* | 12.5 | 8.1 | 0.65 |
| *MEL* | 6.9 | 6.8 | 0.98 |
| *FCA* | n.a. | 0.6 | 0.4* |
| *SJO* | 3.4 | 1.4 | 0.4 |
| *SCB* | n.a. | 1.6 | 0.4* |

*Estimated based on correlation with $Fe^{2+}$ concentration and linear regression within the high-NO group (Fig. S6).
